# HectoSTAR microLED optoelectrodes for large-scale, high-precision *in invo* opto-electrophysiology

**DOI:** 10.1101/2020.10.09.334227

**Authors:** Kanghwan Kim, Mihály Vöröslakos, Antonio Fernández-Ruiz, Saman S. Parizi, Eunah Ko, Blake Hendrix, John P. Seymour, Kensall D. Wise, György Buzsáki, Euisik Yoon

## Abstract

We present a device that can be utilized for a large-scale in vivo extracellular recording, from more than 250 electrodes, with the capability to optically modulate activities of neurons located at more than a hundred individual stimulation targets at the anatomical resolution.

## Introduction

Neurons talk with one another using pulses of electricity. The ability to accurately monitor the change in each neuron’s electrical potential, therefore, enables the study of complicated circuits inside the brain. Not only are the neuronal circuits in the brain densely populated, but also the connections among them exist throughout different parts of the brain [1]. A number of devices that enable highly multiplexed recording of the neurons inside a brain region with single insertion [2-6] have been developed to help neuroscientists answer some of the long-sought-for questions about brain circuits.

The complete understanding of a system’s behavior originates from not the sole knowledge of the connections among the components inside the system but the knowledge combined with the identification of the role of each individual component. Likewise, since the sole observation of the spontaneous activities of the neurons cannot provide sufficient information about the functional connectivity inside the circuit, tools that can provide the capability to precisely modulate a subpopulation of the recorded neuron have been long sought for. A number of devices that enable bidirectional interfacing to the neurons have been developed, and most of them utilize light to control neurons. Optogenetics [7] has been the primary choice to accompany electrical recording since it enables high-resolution, cell-type controlling capability and does not prevent concurrent electrical recording during stimulation [8, 9]. Most devices for the bi-directional opto-electronic neural interfacing (or opto-electrophysiology in short) utilize waveguide structures [10-22] to deliver light generated from an external source to the stimulation site. Because the waveguide occupies a significant amount of space on the surface of the device, however, the approach of steering light from the outside by means of multiple waveguides is not optimal for the precise, multi-site optical stimulation targeting the volume covered by the recording electrodes.

A unique device that enables the multi-site, colocalized light delivery to the very monitored volume [23] was introduced in 2015. An approach to monolithically integrated light sources (LEDs) and the electrodes on the same small platform was first adopted on the device, which is also known as microLED optoelectrode. On a microLED optoelectode, multiple neuron-sized LEDs are placed right next to the electrodes, with less than twenty micrometers of distance in between. The device enabled, for the first time, the high-resolution (60 μm) optical stimulation within a small brain region while the activities of the neurons within the region are recorded [23]. It was the monolithic integration approach that enabled the high-density integration of both the LEDs and the electrodes within a small form factor and that suggested the possibility of large-scale, high-resolution opto-electrophysiology.

Here, we present a device that further extends the capability of the first microLED optoelectrode. The device can provide optical stimuli to more than one hundred stimulation sites located amidst its neuronal signal recording electrodes and thus enables targeting of neurons at the anatomical resolution. To emphasize the device’s capability to deliver optical stimulation to more than hundred (hecto-) stimulation targets at the anatomical resolution, we named the device hectoSTAR microLED optoelectrode. We demonstrated that the hectoSTAR microLED optoelectrode can enable the analysis of wide-ranging, high-density brain circuits with single insertion. Details of the design and the fabrication process of the hectoSTAR microLED optoelectrode is presented. The results of an in vivo experiment performed with a hectoSTAR microLED optoelectrode is presented as well.

## Results

### Design and fabrication of hectoSTAR microLED optoelectrode

The new optoelectrode was designed according to a few guidelines that can maximize the utility of the device. First, the optoelectrode was required to record from as many neurons as possible from a wide brain area. Second, the activity of a neuron within the recorded region should be recorded from preferably more than one electrode. A hypothetical neuronal layer within the region should be able to be selectively stimulated from either above the layer, within the layer, or below the layer using an LED, without having to move the optoelectrode. Finally, the cross-sectional area of each shank of the optoelectrode should be made as small as possible in order to minimize the acute damage induced in the tissue during insertion.

The hectoSTAR microLED optoelectrode was fabricated in a four-shank configuration, where 64 electrodes and 32 LEDs are placed on each of the shanks. A considerably large section of a brain (either coronal or sagittal, depending on the application), whose area is as large as 1.17 mm^2^ (900 × 1,300 μm), can be studied in one insertion of the optoelectrode (Fig. 1A). Two columns of the electrodes, each of which contains thirty-two small (11 × 15 μm) electrodes, were placed at the center of the shank. The vertical distance between two adjacent electrodes on each column and the horizontal distance between the columns were chosen to be 40 μm and 27 μm, respectively, so that the distance between any two adjacent electrodes is no larger than 40 μm. The tight electrode configuration allows multiple electrodes to simultaneously pick up the activities of a neuron and assist spike sorting. The microLEDs (8 × 15 μm) are located at the center of the shank, allowing a precise, co-localized optical stimulation of the neurons whose activities are being monitored by the electrodes. The vertical distance between two adjacent LEDs was set as 40 μm, identical to the vertical pitch of the electrodes on each column.

**Fig 1.**
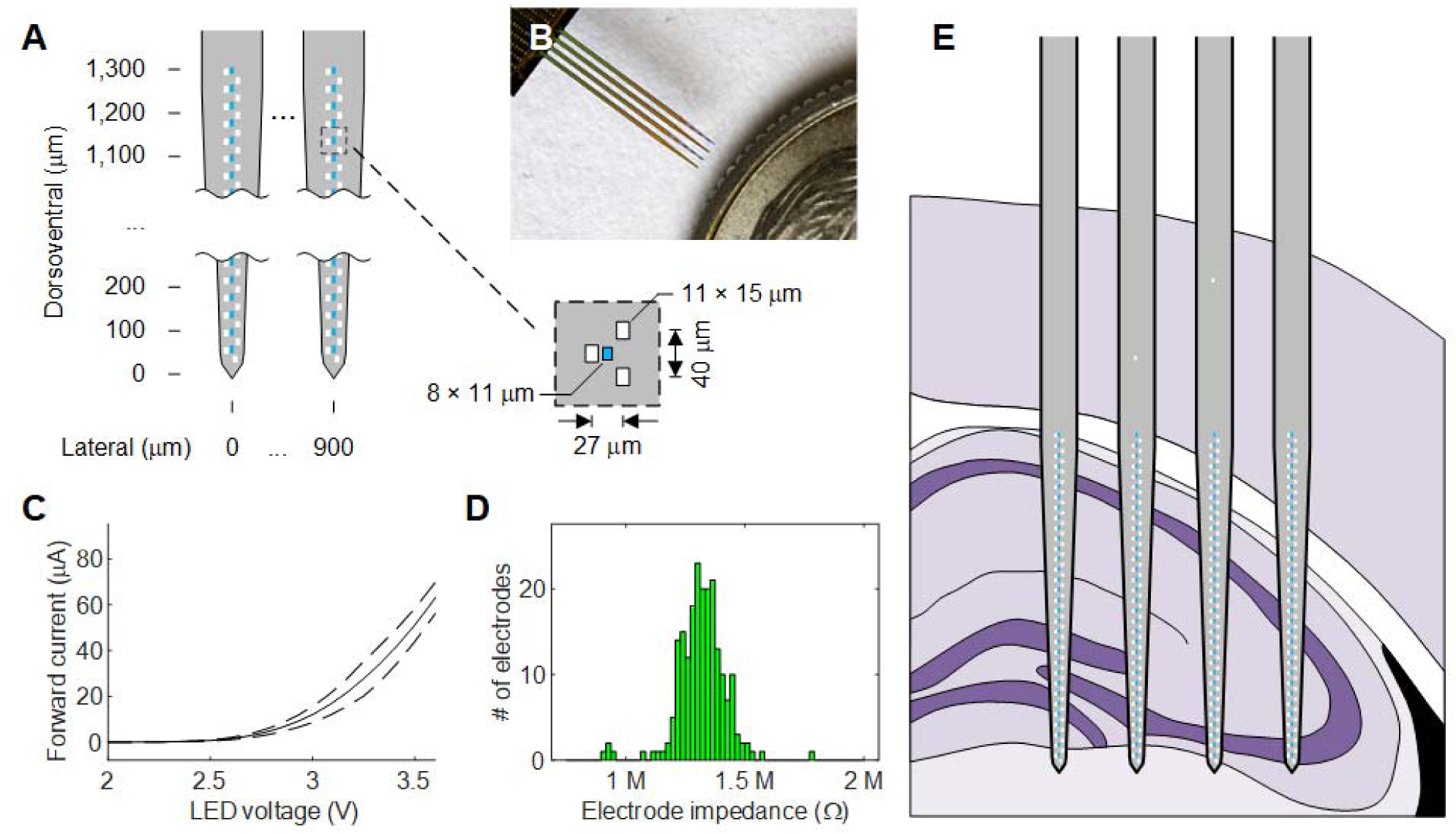
Details of hectoSTAR microLED optoelectrode. **A** A schematic diagram of the tips of a hectoSTAR microLED optoelectrode. Grey polygon indicates silicon shank body, white rectangles electrodes, and light blue rectangles microLEDs. The inset shows the dimensions of the components and the distances between each component. **B** A microphotograph of a fabricated hectoSTAR microLED optoelectrode. **C** The voltage-current relationship of the LEDs on a hectoSTAR microLED optoelectrode. The solid line indicates the mean, and the dashed lines one standard deviation away from the mean. **D** The distribution of the 1-kHz impedance of the electrodes o a hectoSTAR microLED optoelectrode. The mean and the standard deviation of the impedance shown on the histogram are 1.32 and 0.08 MΩ, respectively. **E** The locations of the electrodes and the LEDs on a hypothetical hectoSTAR optoelectrode implanted inside the dorsal hippocampus of a mouse brain. The solid purple line indicates the pyramidal layer in the hippocampus.

The hectoSTAR microLED optoelectrodes were fabricated using miniSTAR technology [24]. GaN-on-Si GaN/InGaN MQW LED wafers with high boron doping density inside the silicon substrate were used, and the signal interconnects were formed in a three-metal-layer configuration [24]. The metallic interconnecting traces (i.e. interconnects) for both the LED drive signal and the recorded neural signals were formed at 700 lines per millimeter density (0.7 μm half-pitch), utilizing a thin lift-off resist and an i-line step-and-repeat projection photolithography tool with 5 × image reduction capability. Figure 1B shows a microphotograph of a fabricated hectoSTAR microLED optoelectrode.

The electrical characteristics of the microLEDs as well as those of the electrodes are presented in Fig. 1C and 1D. The microLEDs turn on (allows 1 μA of current) at 2.55 ± 0.04 V (mean ± SD, n = 128) and allow 51.3 ± 7.4 μA of current (mean ± SD, n = 128) when biased at 3.5 V. Among 256 electrodes on an optoelectrode, 201 electrodes had impedance between 1 MOhms and 1.5 MOhms at 1 kHz.

The large area the electrodes and the LEDs cover allows even a complicated study to be conducted within a single insertion. The vertical and the horizontal coverage is large enough to cover the whole CA3 and a part of CA1 in a mouse’s dorsal hippocampus (Fig. 1E). For example, the signal propagation and the dynamics of long-term potentiation via Schaffer’s collateral can be studied with a single insertion.

### Large-scale recording of spontaneous neuronal activities

We inserted a hectoSTAR optoelectrode into the dorsal hippocampus on the left hemisphere of a mouse’s brain and recorded spontaneous brain activities. A transgenic (Thy1-ChR2-YFP) adult male mouse (28 g) was used. The mouse was head-fixed on a stereotaxic frame and anesthetized with 1.2 % isoflurane inhalant. After a craniotomy (at AP = −2 mm and ML = 1.5 mm, from bregma), the probe was implanted into the brain and slowly lowered until the tips were lowered by approximately 1.5 mm from the brain surface. The approximate location of the probe in the brain is shown in Fig. 2A.

**Fig 2.**
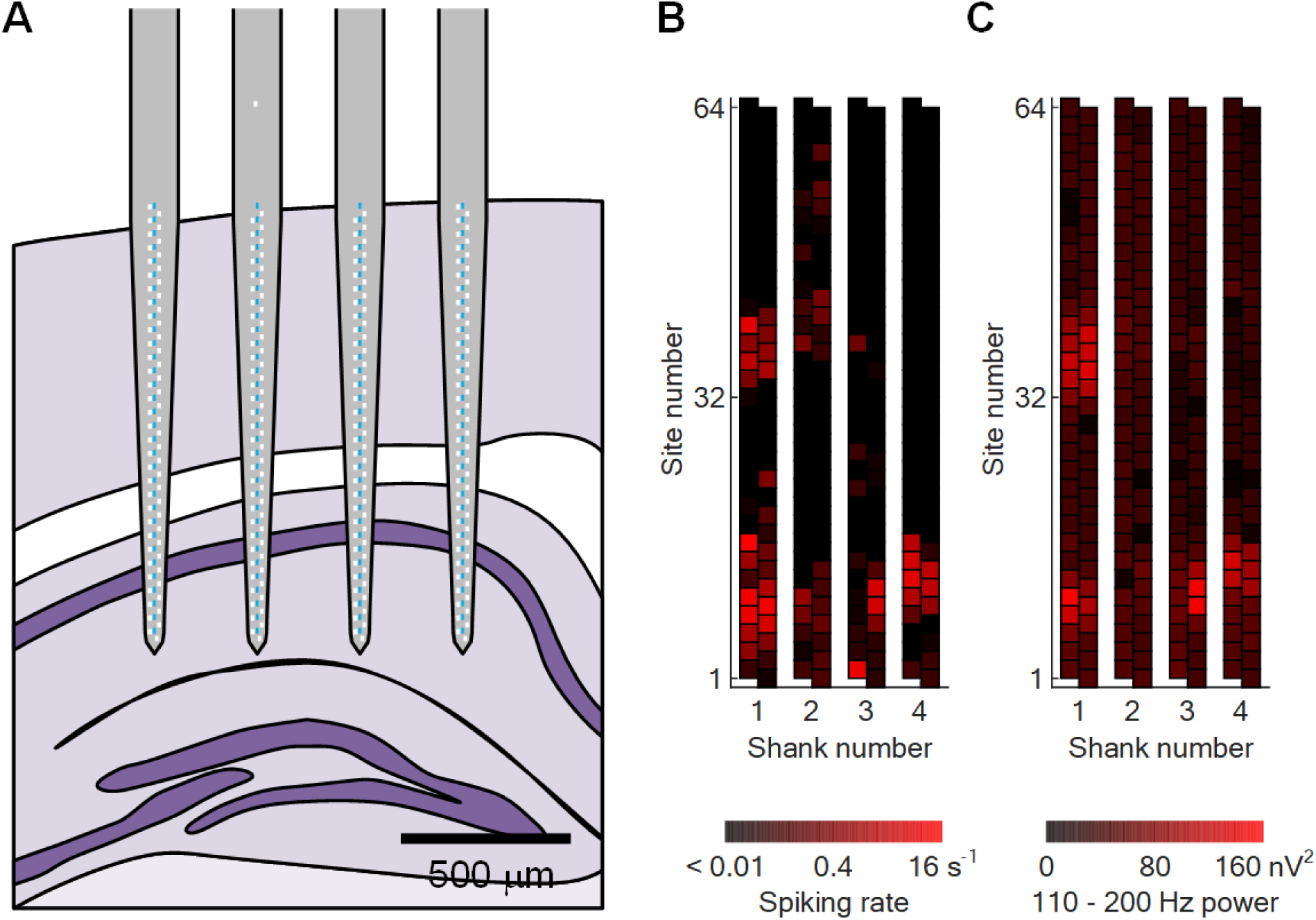
Large-scale recording of spontaneous activities within a mouse brain. **A** The location of the hectoSTAR optoelectrode in the mouse brain. The solid purple layer indicates the hippocampal pyramidal layer. **B** Heatmap of activities with an amplitude larger than 100 μVpp, after high-pass filtering of the signal at 250 Hz, detected from each site. **C** Heatmap of the distribution of fast gamma band (110 - 200 Hz) power in the signal recorded from each electrode. For both heatmaps, a one-minute recording, acquired 15 minutes after the completion of optoelectrode implantation, was utilized for calculation of both values. Each square in each heatmap represents a single electrode.

After the brain recovered for approximately 15 minutes, the spontaneous activities of the neurons in the vicinity of the electrodes were recorded. Wideband (0.1 - 10 kHz) signals were recorded from all the channels, and the recorded signals were analyzed upon the completion of the in vivo experiment.

The variance of the shape and the amplitude of the signals recorded from each electrode on the optoelectrode allowed an easy on-line determination of the accurate positions of the electrodes (as well as the LEDs) on the optoelectrode in the brain. The regions with higher density of somata were clearly highlighted with occurrences of high-amplitude and high-frequency signals, suggestive of sharp waves and ripples (SWRs). Post-hoc analysis of the recorded signals showed that the locations of the electrodes that recorded the most high-amplitude, high-frequency signals (> 100 μV peak-to-peak when high-pass filtered at 250 Hz, shown in Fig. 2B) and those with the highest power of the mid-frequency range signal (‘fast gamma,’ 110 - 200 Hz band, shown in Fig. 2C) corresponds well with one another. From the coordinates of the tips of the optoelectrode, the layers of the somata were identified as a cortical layer (top) and the hippocampal CA1 pyramidal layer (bottom), respectively.

### Precise optical modulation of local neuronal activities

We demonstrated that the microLEDs on the hectoSTAR optoelectrode are capable of inducing a temporally and spatially precise perturbation of the brain circuit. We utilized an LED whose location corresponded to the focus of the activities, i.e. near the electrodes from which activities with the largest amplitude were recorded, near where the hippocampal pyramidal layer is expected to be located (6th LED from the bottom, on the leftmost shank, i.e. shank 1; Fig. 3B). Optical pulses with varying intensities were delivered until induced high-frequency oscillations (iHFOs) were observed in the recordings from the electrodes located in the vicinity of the LED. Voltage pulses with V_on = 3.5 V induced strong iHFOs. Total 260 pulses were generated at 0.2 Hz (t_on = 100 ms).

**Fig 3.**
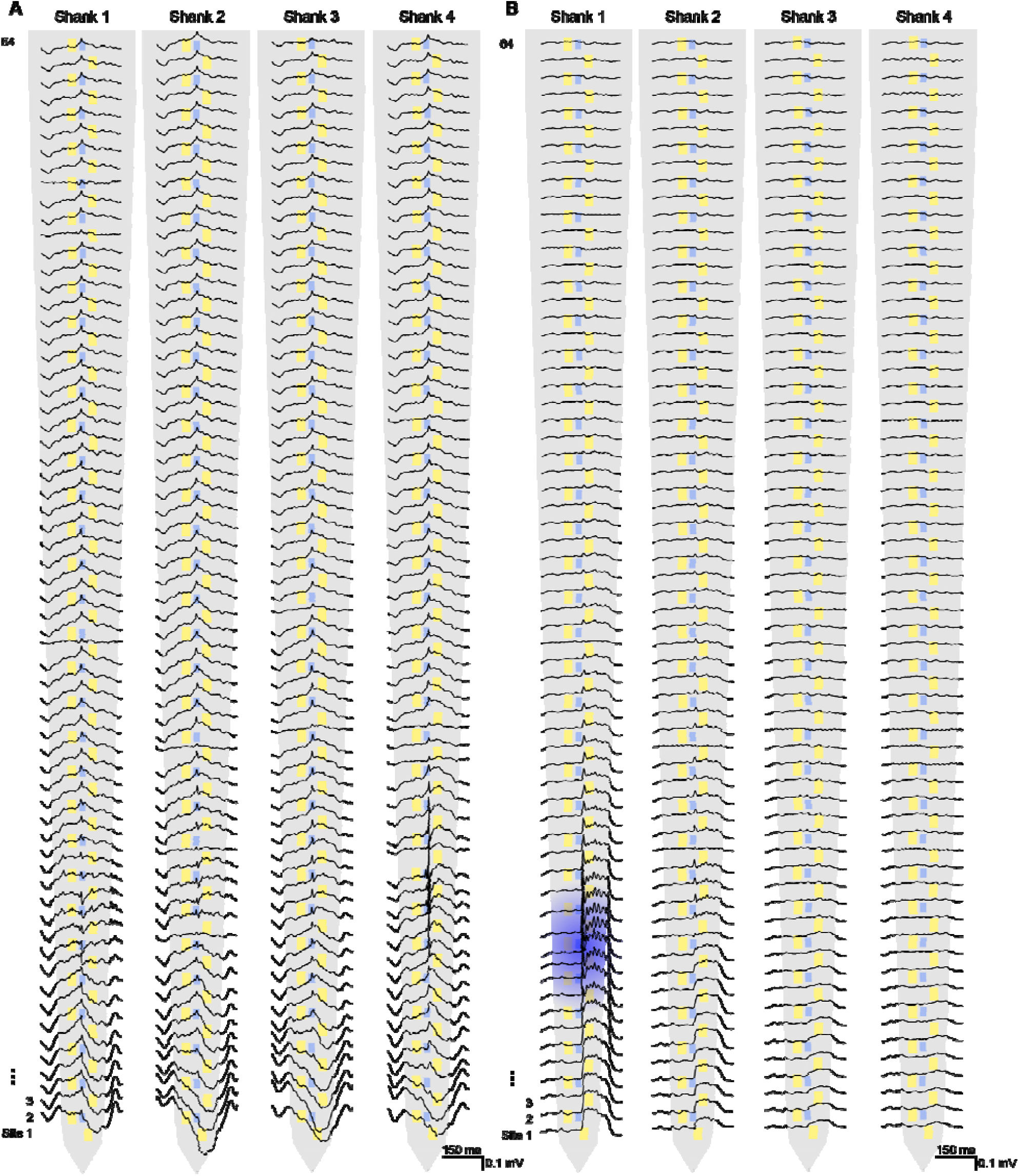
Spatially confined optical stimulation in the hippocampal pyramidal layer. **A** SPW-R triggered, band-pass filtered average waveforms across all the channels (80-250 Hz band-pass, n = 190 SPW-Rs). **B** microLED stimulation triggered, band-pass filtered average waveforms across all the channels. Note, the stimulation-evoked local, high-frequency oscillations in the CA1 region of the hippocampus (100 ms square pulses were delivered on the 6th LED from the tip of shank-1, highlighted in blue; n = 260 pulses at 3.5 V intensity).

**Fig 4.**
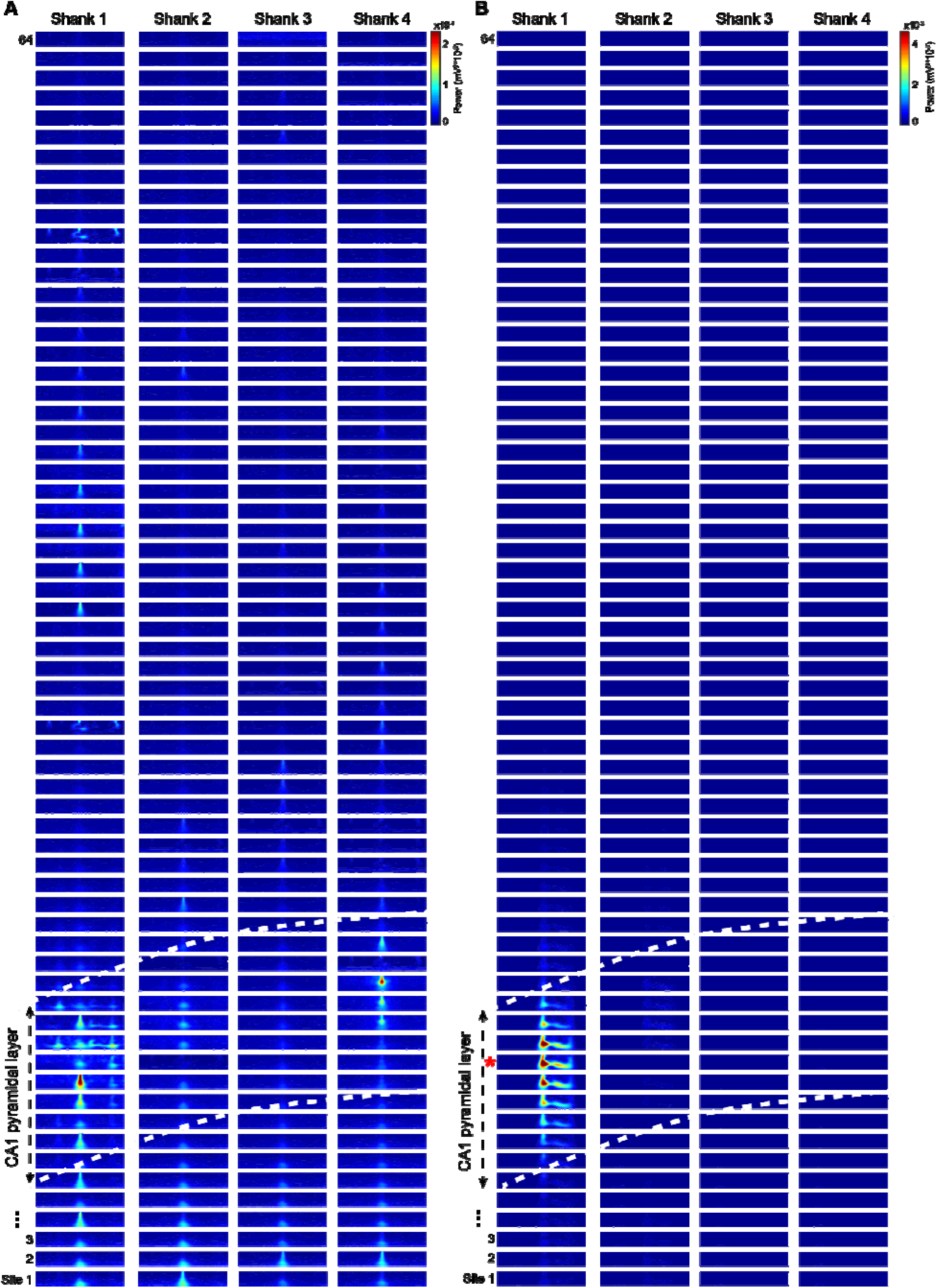
Spatial distribution of spontaneous and induced high-frequency oscillations. **A** Wavelet spectrograms for spontaneous hippocampal sharp-wave ripples (SPW-Rs). Each panel corresponds to one electrode, arranged from most superficial to deepest, with each column representing one each four shanks of the probe. White dashed lines delineate the hippocampal CA1 pyramidal layer. Above it is the neocortex and below CA1 dendritic layers. Each spectrogram shows the power in the 50-150 Hz frequency band in a window of 300 ms around the SPW-Rs detected in the middle of the pyramidal layer in the leftmost shank (n = 190 SPW-Rs). Note that the characteristic high-frequency power of SPW-Rs is present also in the rightmost shank, 900um away from where they were detected, indicating that SPW-Rs are synchronous across the whole CA1 covered by the probe. Power in all panels was normalized to the maximum value across all 256 electrodes to facilitate comparison. **B** The same spectrograms were calculated around the onset of 100ms square pulse stimulations delivered through a microLED located on shank-1 (6^th^ LED from the tip of the electrode; red asterisk) (n = 260 pulses, 3.5V). Note that the stimulation induced a high-frequency oscillation that resemble the spontaneous SPW-Rs, but in this case, it was restricted to ∼ 8 electrodes (160 μm) around the stimulation microLED.

Offline signal analysis revealed that the ripple-like high-frequency oscillation induced by the optical stimulation was strictly confined to a part of the hippocampal pyramidal layer where the illumination was provided. As can be seen on Fig 3, spontaneous sharp wave-ripple complexes (SPW-Rs) were recorded from electrodes located throughout the CA1 region of the hippocampus, with synchronous high-power activity detected in the pyramidal layer (Fig 3A). On the other hand, iHFOs did not propagate from the stimulation site and were recorded only from the electrode located within an 80-um radius from the LED from which the stimulating light was generated (Fig 3B). This result suggests that the light was confined within a very small volume and that only the neurons within the volume contributed to the spatiotemporally confined activity.

### Study of wide-ranging neuronal connections with large-scale, high-resolution opto-electrophysiology

In order to demonstrate hectoSTAR optoelectrode’s capability of the pin-point stimulation of areas within a long-ranging circuit, we provided optical stimulation to another location in the region where the neuronal activities were being recorded. After the optical stimulation of the hippocampal CA1 region was completed, we provided light to a point within the neocortex. The 19th LED (counted from the bottom of the shank) on shank 1 was utilized, as it was located at the center of the region where the most large-amplitude, high-frequency signals, suggestive of action potentials, were recorded (Figures 2B and 2C). Light pulses with a duration and a duty cycle (0.2 Hz, DR = 0.2) identical to those used for the hippocampal stimulation were provided. Local iHFO was observed within the region (data not shown) during the stimulation, and the recorded traces were saved for off-line data analysis.

We identified putative single-unit activities from the recorded signals. We utilized data obtained from all three sessions, namely spontaneous SPW-R recording session, hippocampal optical stimulation session, and cortical optical stimulation session. Spike sorting identified 67 units, whose maximum amplitude activities were recorded from 42 of 256 electrodes on the optoelectrode. As can be seen in Fig. 5B, these electrodes were located throughout the regions from which high-amplitude local field potentials were recorded (as shown in Fig. 2C), with those with the greatest number of detected neurons located at the CA1 pyramidal layer. The change of the firing rate during the optical stimulus, or the gain, was calculated for each identified unit. As expected, neurons with higher gain were observed near the stimulation sites, as can be seen in Fig. 5C (hippocampal stimulation) and 5D (cortical stimulation).

**Fig 5.**
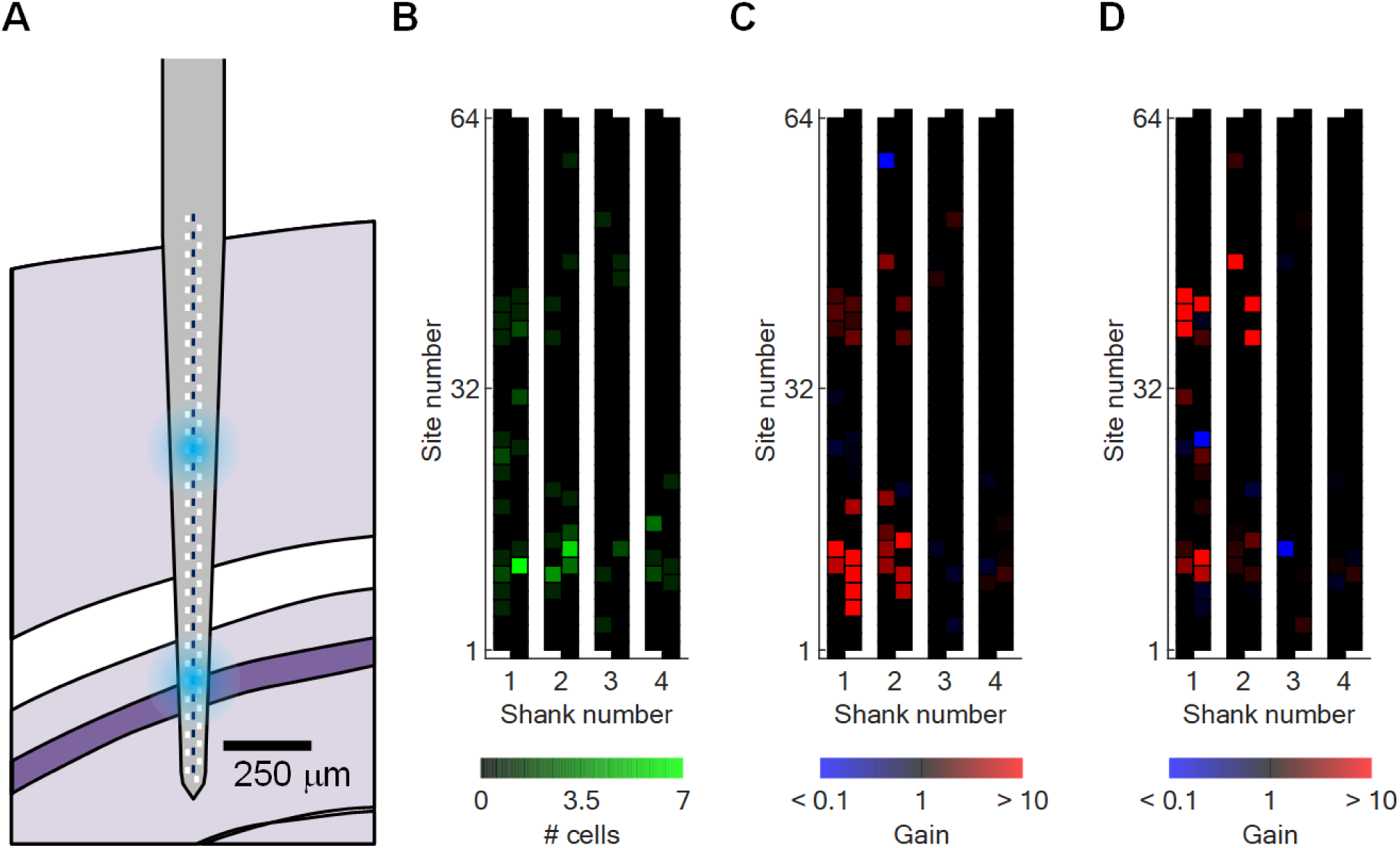
Individual neuronal activity modulation following local optical stimulation. **A** The location of the hectoSTAR optoelectrode in the mouse brain. Blue circles indicate the locations of the microLEDs through which optical stimuli were provided. **B** The number of putative single units whose recorded action potential amplitude was the largest at each electrode. **C** Mean gain of firing rates of neurons whose locations are indicated in B during hippocampal optical stimulation. **D** Mean gain of firing rates of the same neurons during cortical optical stimulation. Each square in each heatmap represents a single electrode.

Further analysis of the firing rate gain of each neuron revealed some interesting patterns that might suggest some unique connections among and within the circuits that exist inside the target region. In general, As shown in Fig. 5C and 5D, neurons near the stimulation sites showed higher gain than those located further away from the sites (Fig. 6B). A closer look at the change of the gain of the individual neurons, however, reveals some interesting behavior of a few of the neurons (Fig. 6C). Among these are neurons being inhibited during local optical stimulation and excited during distal stimulation: one neuron whose largest-amplitude activity was recorded in CA1 showed a gain of 0.01 during hippocampal stimulation, but showed a gain of 10.2 during cortical stimulation; one cortical neuron showed the change of the gain from 1.59 (Hippocampal) to 0.76 (Cortical). Figures 6D and 6E shows peristimulus time histograms (PSTHs) of a few selected neurons whose gains changed the most dramatically between the stimulations from different locations.

**Fig 6.**
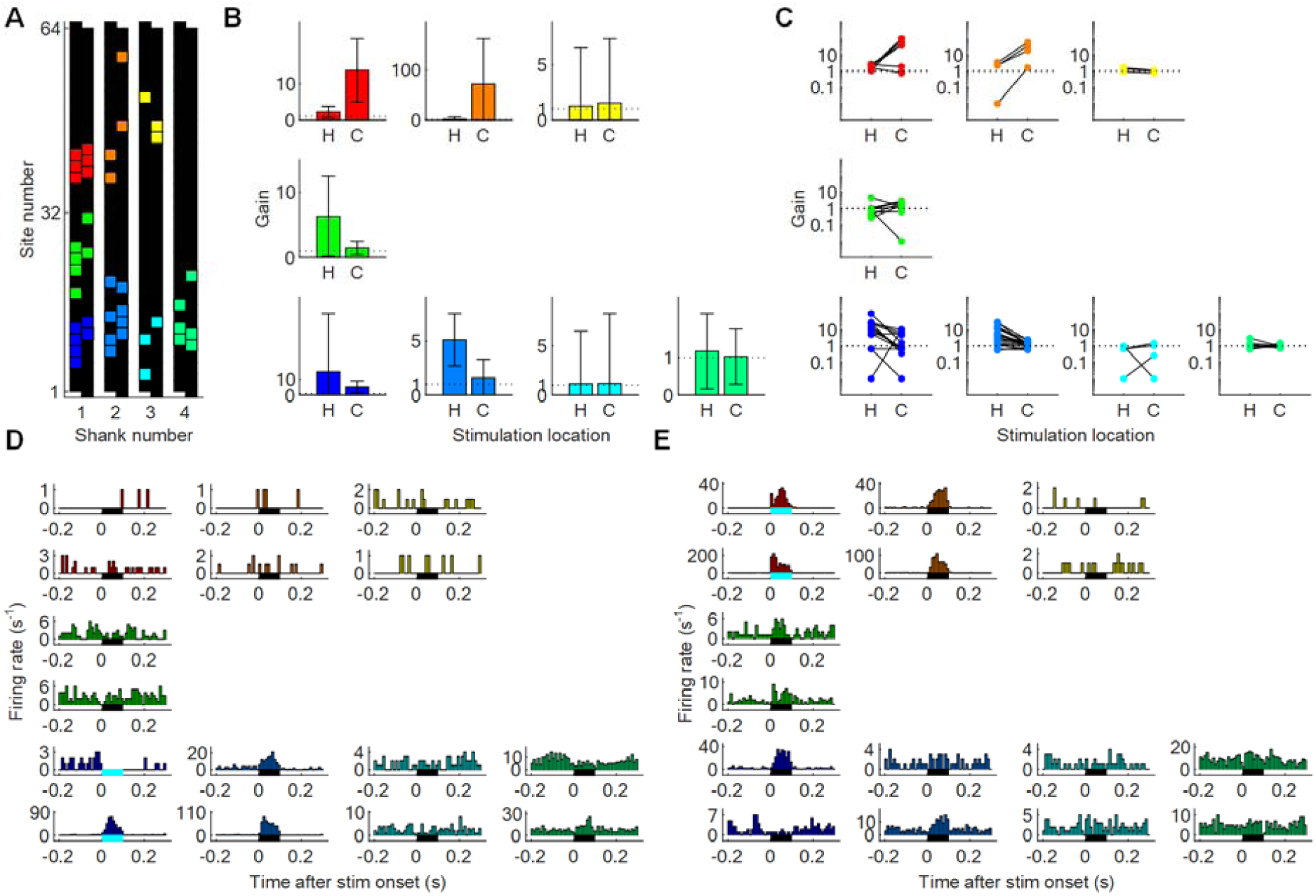
Individual neuronal responses suggesting contributions from both local and long-ranging circuits. **A** Locations of groups of neurons whose collective and individual activity patterns are shown in parts B through E. Neurons whose action potentials were recorded the largest from an electrode are indicated with the same color as the electrode. **B** Firing rate gain of each neuron in each neuronal group whose location is shown in part A during hippocampal and cortical optical stimulation. Boxes indicate the mean and error bars indicate one standard deviation. Patterns similar to those shown in Fig 5C and 5D are visible. **C** Change in the firing rate gain of each neuron in each neuronal group. Closer analysis reveals responses of a few individual neurons acting differently from the other neurons in the same group. **D-E** Peristimulus time histograms (PSTHs) of a few selected neurons from each group during (D) hippocampal stimulation and (E) cortical stimulation. The solid line underneath each histogram indicates the duration (with its length) and the location (with its color; light blue color indicates the presence of the LED in the vicinity) of the optical stimulation. The activity of a hippocampal neuron (whose PSTH is shown on both part D and E on the first column from the left, the second row from the bottom) is inhibited during hippocampal optical stimulation and strongly enhanced during cortical stimulation, suggesting both an inhibitory intra-hippocampal connection and an excitatory cortical-hippocampal connection.

## Discussion

We fabricated hectoSTAR microLED optoelectrodes, microLED optoelectrodes that can be utilized for in vivo experiments for the study of large neuronal circuits. Two hundred and fifty six electrodes and 128 LEDs, distributed on four shanks and covering a large volume with a cross-sectional area of 900 micrometers by 1,300 micrometers, enables a simultaneous recording of neurons distributed inside a large volume while the activity of only few of them can be selectively perturbed at a very high (< 40 μm) spatial resolution. The hectoSTAR microLED demonstrated its capability of high-precision opto-electrophysiology within a mouse brain.

The utility of the hectoSTAR microLED optoelectrodes can further be improved with miniaturization of the device package. We demonstrated the capabilities of the hectoSTAR microLED optoelectrode with a head-fixed animal due to the large size of the packaged device. The size of an unpackaged device is quite small, measuring 4.2 × 11.1 × 0.03 mm (W x L x T), including the backend for the external connection. However, since the printed circuit board, on which the connectors for the interface with the recording and the stimulation system are integrated, had to be designed with 2-mil (0.051 mm) half-pitch, the size of the package device is as large as 60 × 56 x 1 mm (W x L x T). It is expected that, if miniature-sized interface circuit(s) with wire bonding pads in appropriate dimensions and layouts can be integrated with the circuit(s) by the means of a flexible cable (i.e. using microflex technology [25]), the size of the packaged device can be greatly reduced so that the packaged device can be easily mounted on the head of a mouse to allow the animal to freely move during an experiment.

## Methods

### HectoSTAR microLED optoelectrode fabrication and packaging

The microfabrication processes were carried out in Lurie Nanofabrication Facility, University of Michigan, Ann Arbor, MI, USA. Devices were fabricated utilizing steps identical to the steps described in Kim et al. [24], except for a special photolithography procedure utilized to define narrow metal lines (700 nm half-pitch) on the surface of optoelectrode. Narrow metal lines were utilized for both the LED signal interconnects and recorded signal interconnects, located on the first metal layer and the top metal layer, respectively.

Microflex interconnection technique [25] was utilized to provide electrical connections from the outside to the interconnects on the optoelectrode. A polyimide cable with embedded metal lines was fabricated on a silicon wafer and then released from the wafer. Gold ball bumps were formed on both ends of the metal lines to form vertical connections from the metal lines on the cable to the pads located underneath. A K&S ball bonder was utilized for the process.

### Animal experiment

The animal procedure was approved by the Institution Animal Care and Use Committee of the University of Michigan (protocol number PRO-7275). One male transgenic mouse (JAX stock #007612) was utilized for the experiment.

Electrophysiology recordings were made using two RHD 128-channel recording headstages (Intan technologies, Los Angeles, CA) connected to the PCB on which the probe was mounted via two pairs of Molex SlimStack (502426-6410, Molex, Lisle, IL) connectors. A PC running Intan data acquisition software, connected to an Intan USB interface board via a USB 2.0 cable, was utilized to acquire and save data in real-time. NeuroScope [26] was utilized for the real-time visualization of data collected from all the 256 channels.

Voltage signals for the LED driving were provided using a function generator (33220A, Keysight Technologies, Santa Rosa, CA). Rectangular voltage pulses with 0 V low-level voltage and 3.5 V high-level voltage were used as the driving signal, and one-hundred-millisecond-long pulses were applied every 5 seconds (n = 260 pulses).

### Local field potential analysis

Ripple detection and wavelet spectrogram calculation were performed as previously described [27, 28].

To detect ripples a single electrode in the middle of the pyramidal layer was selected. The wide-band LFP signal was band-pass filtered (difference-of-Gaussians; zero-lag, linear phase FIR), and instantaneous power was computed by clipping at 4 SD, rectified and low-pass filtered. The low-pass filter cut-off was at a frequency corresponding to *p* cycles of the mean band-pass (for 80-250 Hz band-pass, the low-pass was 55 Hz). Subsequently, the power of the non-clipped signal was computed, and all events exceeding 4 SD from the mean were detected. Events were then expanded until the (non-clipped) power fell below 1 SD; short events (< 15 ms) were discarded.

To analyze high-frequency oscillatory activity in the LFP at a high resolution in time and frequency, the complex wavelet transform of the LFP was calculated using complex Morlet wavelets. Wavelets were calculated for every 2 Hz frequency step in the 50-150 Hz band. Spectrograms were calculated for each detected SPW-R or stimulation pulse in a [-150, +150] ms window using the LFP from every individual electrode. Spectrograms for individual events were averaged to construct final plots.

### Action potential analysis

A concatenated signal file was prepared by merging all three recordings. Putative single units were first sorted using Kilosort [29] and then manually curated using Phy (https://phy-contrib.readthedocs.io/). After extracting timestamps of each putative single unit activity, peristimulus time histograms and firing rate gains were analyzed using a custom MATLAB (Mathworks, Natick, MA) script.

## Author Information

### Author contributions

K. K., G. B., and E. Y. together worked on the conceptual design of hectoSTAR LED
optoelectrodes. S. S. P. developed the high-density metal patterning process with J. P. S.. K. K. designed hectoSTAR LED optoelectrodes with inputs from M. V., G. B., J. P. S., and E. Y.; fabricated the optoelectrodes; and assembled the optoelectrodes with E. K. and B. H.. B. H. characterized the packaged optoelectrodes with K. K.. M. V. carried out in vivo experiments with K. K. and processed recorded neuronal signals. E. K. took device photographs. K. K., M. V., and E. Y. wrote the manuscript. All authors discussed the results and commented and edited the manuscript.

## References

[1] S. W. Oh et al., “A mesoscale connectome of the mouse brain,” Nature, vol. 508, no. 7495, pp. 207–214, 2014/04/01 2014.

[2] J. Csicsvari et al., “Massively Parallel Recording of Unit and Local Field Potentials With Silicon-Based Electrodes,” Journal of Neurophysiology, vol. 90, no. 2, pp. 1314–1323, 2003/08/01 2003.

[3] R. Fiáth et al., “A silicon-based neural probe with densely-packed low-impedance titanium nitride microelectrodes for ultrahigh-resolution in vivo recordings,” Biosensors and Bioelectronics, vol. 106, pp. 86–92, 2018/05/30/ 2018.

[4] J. J. Jun et al., “Fully integrated silicon probes for high-density recording of neural activity,” Nature, vol. 551, no. 7679, pp. 232–236, 2017/11/01 2017.

[5] M. E. Merriam, S. Dehmel, O. Srivannavit, S. E. Shore, and K. D. Wise, “A 3-D 160-Site Microelectrode Array for Cochlear Nucleus Mapping,” IEEE Transactions on Biomedical Engineering, vol. 58, no. 2, pp. 397–403, 2011.

[6] J. Scholvin et al., “Close-Packed Silicon Microelectrodes for Scalable Spatially Oversampled Neural Recording,” IEEE Transactions on Biomedical Engineering, vol. 63, no. 1, pp. 120–130, 2016.

[7] E. S. Boyden, F. Zhang, E. Bamberg, G. Nagel, and K. Deisseroth, “Millisecond-timescale, genetically targeted optical control of neural activity,” Nature Neuroscience, vol. 8, no. 9, pp. 1263–1268, 2005/09/01 2005.

[8] G. Buzsáki et al., “Tools for Probing Local Circuits: High-Density Silicon Probes Combined with Optogenetics,” Neuron, vol. 86, no. 1, pp. 92–105, 2015/04/08/ 2015.

[9] L. Grosenick, James H. Marshel, and K. Deisseroth, “Closed-Loop and Activity-Guided Optogenetic Control,” Neuron, vol. 86, no. 1, pp. 106–139, 2015/04/08/ 2015.

[10] P. Anikeeva et al., “Optetrode: a multichannel readout for optogenetic control in freely moving mice,” Nature Neuroscience, vol. 15, no. 1, pp. 163–170, 2012/01/01 2012.

[11] A. Canales et al., “Multifunctional fibers for simultaneous optical, electrical and chemical interrogation of neural circuits in vivo,” Nature Biotechnology, vol. 33, no. 3, pp. 277–284, 2015/03/01 2015.

[12] K. Kampasi et al., “Dual color optogenetic control of neural populations using low-noise, multishank optoelectrodes,” Microsystems & Nanoengineering, vol. 4, no. 1, p. 10, 2018/06/04 2018.

[13] K. Kampasi et al., “Fiberless multicolor neural optoelectrode for in vivo circuit analysis,” Scientific Reports, vol. 6, no. 1, p. 30961, 2016/08/03 2016.

[14] Y. LeChasseur et al., “A microprobe for parallel optical and electrical recordings from single neurons in vivo,” Nature Methods, vol. 8, no. 4, pp. 319–325, 2011/04/01 2011.

[15] S. Libbrecht et al., “Proximal and distal modulation of neural activity by spatially confined optogenetic activation with an integrated high-density optoelectrode,” Journal of Neurophysiology, vol. 120, no. 1, pp. 149–161, 2018/07/01 2018.

[16] S. Royer, B. V. Zemelman, M. Barbic, A. Losonczy, G. Buzsáki, and J. C. Magee, “Multi-array silicon probes with integrated optical fibers: light-assisted perturbation and recording of local neural circuits in the behaving animal,” European Journal of Neuroscience, vol. 31, no. 12, pp. 2279–2291, 2010/06/01 2010.

[17] Y. Son et al., “In vivo optical modulation of neural signals using monolithically integrated two-dimensional neural probe arrays,” Scientific Reports, vol. 5, no. 1, p. 15466, 2015/10/23 2015.

[18] E. Stark, T. Koos, and G. Buzsáki, “Diode probes for spatiotemporal optical control of multiple neurons in freely moving animals,” Journal of Neurophysiology, vol. 108, no. 1, pp. 349–363, 2012/07/01 2012.

[19] J. Wang et al., “Integrated device for combined optical neuromodulation and electrical recording for chronicin vivoapplications,” Journal of Neural Engineering, vol. 9, no. 1, p. 016001, 2011/12/07 2011.

[20] L. Wang et al., “An artefact-resist optrode with internal shielding structure for low-noise neural modulation,” Journal of Neural Engineering, vol. 17, no. 4, p. 046024, 2020/08/05 2020.

[21] F. Wu et al., “An implantable neural probe with monolithically integrated dielectric waveguide and recording electrodes for optogenetics applications,” Journal of Neural Engineering, vol. 10, no. 5, p. 056012, 2013/08/28 2013.

[22] J. Zhang et al., “Integrated device for optical stimulation and spatiotemporal electrical recording of neural activity in light-sensitized brain tissue,” Journal of Neural Engineering, vol. 6, no. 5, p. 055007, 2009/09/01 2009.

[23] F. Wu, E. Stark, P.-C. Ku, Kensall D. Wise, G. Buzsáki, and E. Yoon, “Monolithically Integrated μLEDs on Silicon Neural Probes for High-Resolution Optogenetic Studies in Behaving Animals,” Neuron, vol. 88, no. 6, pp. 1136–1148, 2015/12/16/ 2015.

[24] K. Kim, M. Vöröslakos, J. P. Seymour, K. D. Wise, G. Buzsáki, and E. Yoon, “Artifact-free and high-temporal-resolution in vivo opto-electrophysiology with microLED optoelectrodes,” Nature Communications, vol. 11, no. 1, p. 2063, 2020/04/28 2020.

[25] T. Stieglitz, H. Beutel, and J. U. Meyer, ““Microflex”—A New Assembling Technique for Interconnects,” Journal of Intelligent Material Systems and Structures, vol. 11, no. 6, pp. 417–425, 2000/06/01 2000.

[26] L. Hazan, M. Zugaro, and G. Buzsáki, “Klusters, NeuroScope, NDManager: A free software suite for neurophysiological data processing and visualization,” Journal of Neuroscience Methods, vol. 155, no. 2, pp. 207–216, 2006/09/15/ 2006.

[27] A. Fernández-Ruiz, A. Oliva, E. Fermino de Oliveira, F. Rocha-Almeida, D. Tingley, and G. Buzsáki, “Long-duration hippocampal sharp wave ripples improve memory,” Science, vol. 364, no. 6445, p. 1082, 2019.

[28] A. Oliva, A. Fernández-Ruiz, G. Buzsáki, and A. Berényi, “Role of Hippocampal CA2 Region in Triggering Sharp-Wave Ripples,” Neuron, vol. 91, no. 6, pp. 1342–1355, 2016/09/21/ 2016.

[29] M. Pachitariu, N. Steinmetz, S. Kadir, M. Carandini, and H. Kenneth D, “Kilosort: realtime spike-sorting for extracellular electrophysiology with hundreds of channels,” bioRxiv, p. 061481, 2016.

